# Division of Labor in Hand Usage Is Associated with Higher Hand Performance in Free-Ranging Bonnet Macaques, *Macaca radiata*

**DOI:** 10.1101/013268

**Authors:** Madhur Mangalam, Nisarg Desai, Mewa Singh

**Affiliations:** Department of Psychology – University of Georgia, Athens, United States of America; Indian Institute of Science Education and Research Pune, Pune, India; Biopsychology Laboratory, and Institute of Excellence – University of Mysore, Mysore, India; Evolutionary & Organismal Biology Unit – Jawaharlal Nehru Centre for Advanced Scientific Research, Bangalore, India

## Abstract

A practical approach to understanding lateral asymmetries in body, brain, and cognition would be to examine the performance advantages/disadvantages associated with the corresponding functions and behavior. In the present study, we examined whether the division of labor in hand usage, marked by the preferential usage of the two hands across manual operations requiring maneuvering in three-dimensional space (e.g., reaching for food, grooming, and hitting an opponent) and those requiring physical strength (e.g., climbing), is associated with higher hand performance in free-ranging bonnet macaques, *Macaca radiata*. We determined the extent to which the macaques exhibit laterality in hand usage in an experimental unimanual and a bimanual food-reaching task, and the extent to which manual laterality is associated with hand performance in an experimental hand-performance-differentiation task. We observed negative relationships between (a) the latency in food extraction by the preferred hand in the hand-performance-differentiation task (wherein, lower latency implies higher performance), the preferred hand determined using the bimanual food-reaching task, and the normalized difference between the performance of the two hands, and (b) the normalized difference between the performance of the two hands and the absolute difference between the laterality in hand usage in the unimanual and the bimanual food-reaching tasks (wherein, lesser difference implies higher manual specialization). Collectively, these observations demonstrate that the division of labor between the two hands is associated with higher hand performance.

## Introduction

Lateral asymmetries in body, brain, and cognition are almost ubiquitous among biological organisms [1-3]. An adaptationist would advocate that these asymmetries were evolutionarily selected because no bilateral organism can maneuver in three-dimensional space unless one side becomes dominant and always takes the lead [4]. Which side would become dominant, however, is beyond the scope of this hypothesis as there is no advantage or disadvantage evidently associated with either the left or the right side (see Glezer [5], an open-peer commentary on MacNeilage et al. [6]). Among all, manual asymmetries are a central theme of investigation because they are likely to have shaped primate evolution [7]. Manual asymmetries can manifest into (a) hand preference, that is, one hand majorly used while solving a unimanual task (which requires only one hand) or the hand used to execute the most complex action while solving a bimanual task (which requires both hands); (b) hand performance, that is, one hand used to execute actions more efficiently. Fagot and Vauclair [8] reviewed studies on individual- and population-level manual asymmetries among nonhuman primates and proposed the ‘task complexity’ theory which states that the extent of manual asymmetry increases with the complexity of the task (here, the complexity is defined by the spatiotemporal progression of the movements, i.e., coarse verses fine). Observations on several nonhuman primate species are consistent with the task complexity theory. For example, the relatively more complex bimanual food-reaching tasks have been found to elicit greater manual asymmetries than the unimanual versions of the same tasks in capuchin monkeys, *Sapajus* spp. [9,10] and *Cebus capucinus* [11], and chimpanzees, *Pan troglodytes* [12].

Besides exhibiting hand preference and hand performance, several nonhuman primates have been found to exhibit manual specialization, that is, they preferentially use either the left or the right hand while solving some specific types of tasks. For example, while feeding arboreally, captive sifakas, *Propithecus* spp. preferentially used one hand to maintain postural support and the other hand to pluck leaves [13]. While extracting peanut butter from a PVC tube, wild Sichuan snub-nosed monkeys, *Rhinopithecus roxellana* [14], captive tufted capuchin monkeys [15], olive baboons, *Papio anubis* [16], and chimpanzees [17] preferentially used one hand to hold the tube and the other hand to extract the peanut butter. While foraging for food scattered on the ground, captive gorillas, *Gorilla gorilla* [18] and chimpanzees [19] preferentially used one hand to take the food items towards the mouth, and the other hand to hold the remaining ones. While extracting peanuts from a lidded box captive tufted capuchin monkeys consistently used one hand to open the lid of the box and the other hand to reach for them [20]. While allogrooming, wild Sichuan snub-nosed monkeys [21] and both captive and wild chimpanzees [22] preferentially used one hand to hold the skin, and the other hand to remove dirt and ectoparasites. Mangalam et al. [23] argued that these observations might reflect specialization of the two hands for manual actions requiring different dexterity types (i.e., simple/complex hand movements in three-dimensional space, grasping, supporting the body, etc.), and along similar lines described division of labor in hand usage in free-ranging bonnet macaques, *Macaca radiata*. The macaques preferentially used the ‘preferred’ hand for manual actions requiring maneuvering in three-dimensional space (reaching for food, grooming, and hitting an opponent), and the ‘nonpreferred’ hand for those requiring physical strength (climbing). In a hand-performance-differentiation task that ergonomically forced the usage of one particular hand, the macaques extracted food faster with the maneuvering hand compared to the supporting hand, demonstrating the higher maneuvering dexterity of the maneuvering hand. However, whether such division of labor in hand usage improves hand performance in terms of the time and/or energy required to solve a given task remains unexplored.

In the present study, we examined whether the division of labor in hand usage, as described by Mangalam et al. [23], is associated with higher hand performance in free-ranging bonnet macaques, *Macaca radiata*. To this end, we determined the extent to which the macaques exhibit laterality in hand usage in two experimental unimanual and a bimanual food-reaching task, and the extent to which manual laterality is associated with hand performance in an experimental hand-performance-differentiation task. If the division of labor between the two hands is associated with higher hand performance, we would expect negative correlations between (a) the latency in food extraction by the preferred hand in the hand-performance-differentiation task and the normalized difference between the performance of the two hands, which would imply that the macaques that show a greater difference in the performance of the two hands perform better than those that exhibit a smaller difference; and (b) the normalized difference between the performance of the two hands and the absolute difference between the laterality in hand usage in the unimanual and the bimanual food-reaching tasks, which would imply that the macaques that exhibit a higher manual specialization show a greater difference in the performance of the two hands.

## Methods

### Subjects and Study Site

The subjects were 16 free-ranging bonnet macaques: 2 adult males – AM1 and AM2, 1 subadult male – SM1, 4 juvenile males – JM1, JM2, JM3, and JM4, 8 adult females – AF1, AF2, AF3, AF4, AF5, AF6, AF7, and AF8, and 1 juvenile female – JF1 (see Table 1), inhabiting the Chamundi Hill range in Mysore, India (GPS coordinates: 2°14′41″N 76°40′55″E) (Mangalam et al. [23] referred to AF5 as JF1, AF6 as JF2, AF7 as JF3, AF8 as JF4, and JF1 as JF5).

**Table 1.** Raw data on hand usage for the macaques in the unimanual and the bimanual food-reaching tasks (n = 16), and the hand-performance-differentiation task (n = 10).

| Individual | Hand usage in the food-reaching tasks |  |  |  |  |  |  |  | Hand usage in the hand-performance-differentiation task |  |  |
| --- | --- | --- | --- | --- | --- | --- | --- | --- | --- | --- | --- |
|  | Tasks |  |  |  |  |  | Outcomes |  | Latency in food extraction |  | Laterality in hand performance (LHP) |
| | Unimanual (U) | | | Bimanual (B) | | | PH | Abs. ($HI_{Bimanual} - HI_{Unimanual}$) | PH (s) | NPH (s) | |
|  | L | R | HI | L | R | HI |  |  |  |  |  |
| AM1 | 19 | 2 | - 0.810 | 21 | 0 | - 1.000 | L | 0.190 | - | - | - |
| AM2 | 0 | 21 | 1.000 | 0 | 21 | 1.000 | R | 0.000 | - | - | - |
| SM1 | 0 | 21 | 1.000 | 0 | 21 | 1.000 | R | 0.000 | - | - | - |
| JM1 | 0 | 21 | 1.000 | 1 | 20 | 0.905 | R | 0.095 | 2.847 | 3.040 | 0.033 |
| JM2 | 5 | 16 | 0.524 | 21 | 0 | - 1.000 | L | 1.523 | 3.696 | 3.856 | 0.021 |
| JM3 | 21 | 0 | - 1.000 | 21 | 0 | - 1.000 | L | 0.000 | 1.887 | 3.968 | 0.355 |
| JM4 | 20 | 1 | - 0.905 | 21 | 0 | - 1.000 | L | 0.095 | - | - | - |
| AF1 | 1 | 20 | 0.905 | 0 | 21 | 1.000 | R | 0.095 | - | - | - |
| AF2 | 0 | 21 | 1.000 | 0 | 21 | 1.000 | R | 0.000 | 2.440 | 4.360 | 0.282 |
| AF3 | 0 | 21 | 1.000 | 1 | 20 | 0.905 | R | 0.095 | 3.152 | 4.420 | 0.167 |
| AF4 | 21 | 0 | - 1.000 | 18 | 3 | - 0.714 | L | 0.286 | 2.250 | 3.890 | 0.267 |
| AF5 | 15 | 6 | - 0.429 | 21 | 0 | - 1.000 | L | 0.571 | 2.184 | 2.960 | 0.151 |
| AF6 | 20 | 1 | - 0.905 | 21 | 0 | - 1.000 | L | 0.095 | 2.440 | 3.147 | 0.126 |
| AF7 | 15 | 6 | - 0.429 | 20 | 1 | - 0.905 | L | 0.476 | 4.504 | 4.960 | 0.048 |
| AF8 | 13 | 8 | - 0.238 | 6 | 15 | 0.429 | R | 0.666 | - | - | - |
| JF1 | 1 | 20 | 0.905 | 1 | 20 | 0.905 | R | 0.000 | 1.772 | 3.568 | 0.336 |

‘L’ and ‘R’ indicate the usage of left and right hand respectively; PH and NPH indicate the preferred (i.e., maneuvering) and the nonpreferred (i.e., supporting) hand respectively.

### Ethics Statement

We adhered to the American Society of Primatologists (ASP) “Principles for the Ethical Treatment of Nonhuman Primates” and conducted the present study as a part of an ongoing research project that was approved by the Institutional Animal Ethics Committee (IAEC) at the University of Mysore (because we conducted our research on individuals which (a) did not belong to an endangered or a protected species, and (b) inhabited an unprotected land with an unrestricted public access, our research work did not require permission from any other authority).

### Experimental Procedure

We presented the macaques with 3 sets of 7 consecutive trials, that is, 21 trials, of experimental unimanual and bimanual food-reaching tasks. Solving the unimanual task required obtaining a grape from an unlidded wire mesh box (dimensions: 7.5 cm × 7.5 cm × 17.5 cm; these dimensions allowed the usage of only one particular hand at a time) fixed on a wooden platform (dimensions: 90 cm × 60 cm) with one hand (Fig. 1A; Movie S1), whereas solving the bimanual task required opening and supporting the lid of a lidded wire mesh box with one hand and obtaining a grape with the other hand (Fig. 1B; Movie S2). We placed the task apparatus on the ground within ca. 1 m from the focal macaque when no conspecific was present within ca. 3 m from it and observed the corresponding hand usage.

**Figure 1.** **Apparatuses for the unimanual food-reaching task (a), the bimanual food-reaching task (b), and the hand-performance-differentiation task (c).** Reproduced, with permission from Wiley Periodicals, Inc., from Mangalam et al. [23] © 2013 Wiley Periodicals, Inc.

We then presented the macaques with a single trial of an experimental hand-performance-differentiation task that forced the usage of either the left or the right hand. Solving this task required obtaining grapes from the wire mesh boxes attached towards the bottom on the either lateral extremities of a wooden platform (dimensions: 90 cm × 60 cm); this setup ergonomically forced the macaques to use either the left or the right hand (Fig. 1C; Movie S3). We put 7 grapes in one of the boxes, placed the task apparatus on the ground when no conspecific was present within ca. 3 m from the focal macaque, and video recorded the corresponding extraction behavior. We then repeated the same procedure, but this time by putting the grapes in the other box. The macaques mostly took 4 to 7 bouts to take all 7 grapes out of the box. We analyzed the obtained videos frame-by-frame to determine the average latency in food extraction for all the bouts (each bout measured from when the hand entered the box to when it exited) to the nearest 0.04 s.

For each macaque, we determined the handedness index (HI) values for taking the food out of the wire mesh box in the unimanual and the bimanual food-reaching tasks, using the formula: HI = (R − L)/(R + L) (where ‘R’ and ‘L’ represent the frequency of usage of the right and the left hand respectively). The obtained **HI** values could range from − 1 to + 1, with positive values indicating a bias towards the right-hand use and negative values indicating a bias towards the left-hand use, and the absolute **HI** values indicating the strength of the bias. We then determned the absolute difference between the laterality in hand usage in the unimanual and the bimanual food-reaching tasks (lesser difference = higher manual specialization), using the formula = abs. **(HI**_Bimaual_ – **Hl**_Unimanual_). We determined the hand majorly used for taking the food out of the box in the bimanual food-reaching task, which we referred to as the ‘preferred hand,’ and the opposite hand, which we referred to as the ‘nonpreferred hand’ (previously, in Mangalam et al. [23], we referred to these as the ‘maneuvering’ and the ‘supporting’ hand respectively). Moreover, we determined the laterality in hand performance (LHP) in the hand-performance-differentiation task, using the formula: LHP = (latency in food extraction using the nonpreferred hand − latency in food extraction using the preferred hand)/ (latency in food extraction using the nonpreferred hand + latency in food extraction using the preferred hand). The obtained LHP values could range from − 1 to + 1, indicating the normalized difference in the performance between the two hands.

## Results

Table 1 reports the raw data on hand usage for the macaques (whereas all 16 macaques responded to the unimanual and the bimanual food-reaching tasks, only 10 macaques responded to the hand-performance-differentiation task perhaps because of a lower motivation to solve a relatively more difficult and time-consuming activity). We found strong negative correlations between (a) the latency in food extraction by the preferred hand in the hand-performance-differentiation task and the laterality in hand performance (LHP) (Spearman’s rank correlation: r_s_ = − 0.772, n = 10, p = 0.009; Fig. 2A), and (b) the LHP in the hand-performance-differentiation task and the absolute difference between the laterality in hand usage in the unimanual and the bimanual food-reaching tasks (Spearman’s rank correlation: r_s_ = − 0.752, n = 10, p = 0.012; Fig. 2B). There was no difference between the two hands in the number of bouts for taking all 7 grapes out of the box in the hand-performance-differentiation task (Wilcoxon signed-rank test: Z = − 1.511, p = 0.131).

**Figure 2.** **Relationship between (a) the latency in food extraction using the preferred hand (i.e., the maneuvering hand, see mangalam et al. [1]) and the laterality in hand performance (LHP) in the handperformance-differentiation task, and (b) the LHP in the handperformance-differentiation task and the absolute difference between the laterality in hand usage in the unimanual and the bimanual food- reaching tasks, n = 10.**

## Discussion

We examined whether the division of labor in hand usage, as described by Mangalam et al. [23], is associated with higher hand performance in free-ranging bonnet macaques. We observed negative relationships between (a) the latency in food extraction by the preferred hand in the hand-performance-differentiation task (lower latency = higher performance), the preferred hand determined using the bimanual food-reaching task, and the normalized difference between the performance of the two hands, and (b) the normalized difference between the performance of the two hands and the absolute difference between the laterality in hand usage in the unimanual and the bimanual food-reaching tasks (lesser difference = higher manual specialization). These correlations demonstrate that the division of labor between the two hands is associated with higher hand performance: the macaques that exhibit a higher manual specialization show a greater difference in the performance of the two hands, and also perform better than those that exhibit a smaller difference.

On the one hand, the almost ubiquitous existence of manual asymmetries in nonhuman primates is likely to have some ecological advantages, and even more likely when there are underlying neurological asymmetries, as demonstrated in capuchin monkeys [24-27] and chimpanzees [28-30]. On the other hand, there may be some obvious disadvantages. Objects supposedly are randomly located with respect to the midsagittal plane of an individual (i.e., towards the left or towards the right); this introduces difficulty in solving some tasks for individuals having a bias for one particular side. Fagot and Vauclair [8] reviewed studies on manual asymmetries in nonhuman primates and drew a distinction between hand preference and manual specialization. According to them, hand preference refers to the consistent usage of one hand to solve familiar, relatively simple, and highly practiced tasks, and may not be necessarily accompanied by an improvement in hand performance. In contrast, manual specialization refers to the consistent usage of one hand to solve novel, relatively complex, and not-practiced tasks that require peculiar action patterns, and is necessarily accompanied by an improvement in hand performance. Moreover, individuals generally exhibit manual specialization only when the tasks involve cognitively demanding manual actions. Thus, there exists a marked difference between hand preference and manual specialization in terms of the resulting differences in the performance of the two hands, which is evidently visible while considering the forms and/or functions of manual asymmetries, as described by Mangalam et al. [23]. The difference in the HI values between the unimanual and the bimanual food-reaching tasks allowed us quantifying manual specialization as an entity separate from hand preference (which an individual is likely to show because of an inherent bias) and examining whether it is associated with a higher difference in the performance between the two hands.

In a previous study [31], captive capuchin monkeys exhibited a weak, but statistically nonsignificant, positive relationship between the strength of hand preference and the corresponding hand performance in a unimanual and a bimanual versions of the box task. The study acknowledged that the strength of hand preference could have affected the timing of the movements, and so the observed relationship. This was, however, not the case of the present study because the hand-performance-differentiation task ergonomically forced the macaques to use either the left or the right hand, which allowed measuring the hand performance independent of any ceiling effects, i.e., it was unlikely to prime any motor actions associated with the hand opposite to that of the intended one. It provided a standard setup, which could be more widely used to compare hand performance across individuals while minimizing the possibilities of confounding effects. We suggest the development of such standard and robust experimental setups which might help answering the prevailing questions on manual asymmetries in nonhuman primates.

---

**S1 Movie.** This footage illustrates the adult female bonnet macaque – ‘AF5’, solving the unimaunal food-reaching task. Reproduced, with permission from Wiley Periodicals, Inc., from Mangalam et al. [23] © 2013 Wiley Periodicals, Inc.

**S2 Movie.** This footage illustrates the adult female bonnet macaque – ‘AF5’, solving the bimanual food-reaching task. Reproduced, with permission from Wiley Periodicals, Inc., from Mangalam et al. [23] © 2013 Wiley Periodicals, Inc.

**S3 Movie.** This footage illustrates the adult female bonnet macaque – ‘AF5’, solving the hand-performance-differentiation task. Reproduced, with permission from Wiley Periodicals, Inc., from [Mangalam et al. [23]] © 2013 Wiley Periodicals, Inc.

## References

1. Geschwind N, Galaburda AM (1987) Cerebral lateralization: biological mechanisms, associations and pathology. Cambridge, MA: MIT Press.

2. Ward JP, Hopkins WD (1993) Primate Laterality: Current Behavioral Evidence of Primate Asymmetries. New York: Springer-Verlag.

3. Andrew RJ, Rogers LJ (2002) The nature of lateralization in tetrapods. In: Rogers LJ, Andrew RJ, editors. Comparative vertebrate lateralization. Cambridge: Cambridge University Press, pp. 94-125.

4. Holló G, Novák M (2012) The manoeuvrability hypothesis to explain the maintenance of bilateral symmetry in animal evolution. Biol Direct 7: 22. doi: 10.1186/1745-6150-7-22.

5. Glezer II (1987) The riddle of Carlyle: the unsolved problem of the origin of handedness. Behav Brain Sci 10: 273–275. doi.

6. MacNeilage PF, Studdert-Kennedy MJ, Lindblom B (1987) Primate handedness reconsidered. Behav Brain Sci 10: 247-263. doi: 10.1017/SO140525X00047695.

7. Bradshaw JL, Rogers LJ (1993) The evolution of lateral asymmetries, language, tool use, and intellect. San Diego: Academic Press.

8. Fagot J, Vauclair J (1991) Manual laterality in nonhuman primates: a distinction between handedness and manual specialization. Psychol Bull 109: 76-89. doi: 10.1037/0033-2909.109.1.76.

9. Lilak AL, Phillips KA (2008) Consistency of hand preference across low-level and high-level tasks in Capuchin monkeys (*Cebus apella*). Am Journal Primatol 70: 254-260. doi: 10.1002/ajp.20485.

10. Spinozzi G, Castorina MG, Truppa V (1998) Hand preferences in unimanual and coordinated-bimanual tasks by tufted capuchin monkeys (*Cebus apella*). J Comp Psychol 112: 183-191. doi: 10.1037/0735-7036.112.2.183.

11. Meunier H, Vauclair J (2007) Hand preferences on unimanual and bimanual tasks in white-faced capuchins *(Cebus capucinus)*. Am J Primatol 69: 1064-1069. doi: 10.1002/ajp.20437.

12. Hopkins WD, Rabinowitz DM (1997) Manual specialisation and tool use in captive chimpanzees (*Pan troglodytes*): the effect of unimanual and bimanual strategies on hand preference. Laterality 2: 267-277. doi: 10.1080/713754273.

13. Milliken GW, Ferra G, Kraiter KS, Ross CL (2005) Reach and posture hand preferences during arboreal feeding in sifakas (*Propithecus* sp.): a test of the postural origins theory of behavioral lateralization. J Comp Psychol 119: 430-439. doi: 10.1037/0735-7036.119.4.430.

14. Zhao D, Hopkins WD, Li B (2012) Handedness in nature: first evidence on manual laterality on bimanual coordinated tube task in wild primates. Am J Phys Anthropol 148: 36-44. doi: 10.1002/ajpa.22038.

15. Spinozzi G, Lagana T, Truppa V (2007) Hand use by tufted capuchins (*Cebus apella*) to extract a small food item from a tube: digit movements, hand preference, and performance. Am J Primatol 69: 336-352. doi: 10.1002/ajp.20352.

16. Vauclair J, Meguerditchian A, Hopkins WD (2005) Hand preferences for unimanual and coordinated bimanual tasks in baboons (*Papio anubis*). Cogn Brain Res 25: 210-216. doi: 10.1016/j.cogbrainres.2005.05.012.

17. Hopkins WD (1995) Hand preferences for a coordinated bimanual task in 110 chimpanzees (*Pan troglodytes*): cross-sectional analysis. J Comp Psychol 109: 291-297. doi: 10.1037/0735-7036.109.3.291.

18. Meguerditchian A, Calcutt SE, Lonsdorf EV, Ross SR, Hopkins WD (2010) Brief communication: captive gorillas are right-handed for bimanual feeding. Am J Phys Anthropol 141: 638-645. doi: 10.1002/ajpa.21244.

19. Hopkins WD (1994) Hand preferences for bimanual feeding in 140 captive chimpanzees (*Pan troglodytes*): rearing and ontogenetic determinants. Am J Primatol 27: 395-407. doi: 10.1002/dev.420270607.

20. Spinozzi G, Truppa V (2002) Problem-solving strategies and hand preferences for a multicomponent task by tufted capuchins (*Cebus apella*). Int J Primatol 23: 621-638. doi: 10.1023/A:1014977818853.

21. Zhao D, Gao X, Li B (2010) Hand preference for spontaneously unimanual and bimanual coordinated tasks in wild Sichuan snub-nosed monkeys: implication for hemispheric specialization. Behav Brain Res 208: 85-89. doi: 10.1016/j.bbr.2009.11.011.

22. Hopkins WD, Russell JL, Remkus M, Freeman H, Schapiro SJ (2007) Handedness and grooming in *Pan troglodytes*: comparative analysis between findings in captive and wild individuals. Int J Primatol 26: 1315-1326. doi: 10.1007/s10764-007-9221-x.

23. Mangalam M, Desai N, Singh M (2014) Division of labor in hand usage in free-ranging bonnet macaques, *Macaca radiata*. Am J Primatol 76: 576-585. doi: 10.1002/ajp.22250.

24. Phillips KA, Hopkins WD (2007) Exploring the relationship between cerebellar asymmetry and handedness in chimpanzees (*Pan troglodytes*) and capuchins (*Cebus apella*). Neuropsychologia 45: 2333-2339. doi: 10.1016/j.neuropsychologia.2007.02.010.

25. Phillips KA, Sherwood CC (2005) Primary motor cortex asymmetry is correlated with handedness in capuchin monkeys (*Cebus apella*). Behav Neurosci 119: 1701-1704. doi: 10.1037/0735-7044.119.6.1701.

26. Phillips KA, Sherwood CC (2007) Cerebral petalias and their relationship to handedness in capuchin monkeys (*Cebus apella*). Neuropsychologia 45: 2398-2401. doi: 10.1016/j.neuropsychologia.2007.02.021.

27. Phillips KA, Sherwood CC, Lilak AL (2007) Corpus callosum morphology in capuchin monkeys is influenced by sex and handedness. PLOS ONE 2: e792. doi: 10.1371/journal.pone.0000792.

28. Hopkins WD, Cantalupo C (2004) Handedness in chimpanzees (*Pan troglodytes*) is associated with asymmetries of the primary motor cortex but not with homologous language areas. Behav Neurosci 118: 1176-1183. doi: 10.1037/0735-7044.118.6.1176.

29. Hopkins WD, Cantalupo C, Taglialatela J (2007) Handedness is associated with asymmetries in gyrification of the cerebral cortex of chimpanzees. Cereb Cortex 17: 1750–1756. doi: 10.1093/cercor/bhl085.

30. Hopkins WD, Dunham L, Cantalupo C, Taglialatela J (2007) The association between handedness, brain asymmetries, and corpus callosum size in chimpanzees (*Pan troglodytes*). Cereb Cortex 17: 1757-1765. doi: 10.1093/cercor/bhl086.

31. Fragaszy DM, Mitchell SR, 275. (1990) Hand preference and performance on unimanual and bimanual tasks in capuchin monkeys (*Cebus apella*). J Comp Psychol 104: 275-282. doi: 10.1037/0735-7036.104.3.275.

